# Rescue of an Aggressive Female Sexual Courtship in Mice by CRISPR/cas9 Secondary Mutation *in vivo*

**DOI:** 10.1101/226639

**Authors:** Jozsef Zakany, Denis Duboule

**Affiliations:** Department of Genetics and Evolution, University of Geneva, Switzerland; School of Life Sciences, Ecole Polytechnique Fédérale, Lausanne, Switzerland

## Abstract

We had previously reported [1] a mouse line carrying the *Atypical female courtship (HoxD^Afc^*) allele, where an ectopic accumulation of *Hoxd10* transcripts was observed in a sparse population of cells in the adult isocortex, as a result of a partial deletion of the *HoxD* gene cluster (Figure 1A). Female mice carrying this allele displayed an exacerbated paracopulatory behavior, culminating in a severe mutilation of the studs’ external genitals. To unequivocally demonstrate that this intriguing phenotype was indeed caused by an illegitimate function of the HOXD10 protein, we use CRISPR/Cas9 technology to induced a microdeletion into the homeobox of the *Hoxd10* gene in *cis* with the *HoxD^Afc^* allele [2]. Females carrying this novel *HoxD^Del(1-9)d10hd^* allele no longer mutilate males. We conclude that a brain malfunction leading to a severe pathological behavior can be caused by the mere binding to DNA of a transcription factor expressed ectopically. We also show that in *HoxD^Afc^* mice, *Hoxd10* was expressed in cells containing *Gad1* and *Cck* transcripts, corroborating our proposal that a small fraction of GABAergic neurons in adult hippocampus may participate to some aspects of female courtship.

---

Although the heterozygous *HoxD^Afc^* genotype proved semi-lethal in both sexes, only sexually mature females displayed an aberrant courtship behavior. When placed with a male for mating, and regardless of the male genotype (i.e. *HoxD^Afc^* heterozygous or wildtype), females repeatedly bit and injured the male penises, often up to their complete ablation. In such adult *HoxD^Afc^* heterozygous mice, ectopic *Hoxd10* transcript accumulation was found in numerous scattered cells in the hippocampus [1], while *Hox* genes are never expressed rostral to the hindbrain and its derivatives [3]. To confirm the causal role of *Hoxd10* ectopic expression in this unusual behavior, we induced a deletion in the homeobox of the *Hoxd10* gene *in cis* with the *HoxD^Afc^* allele (Figure 1A). Non-homologous end joining of genomic DNA after exposure to a single guide RNA and the Cas9 endonuclease in fertilized eggs resulted in a 10 base-pair large deficiency in the *Hoxd10* homeobox, giving rise to the *HoxD^Del(1-9)d10hd^* allele. This mutant allele had lost the third alpha-helix of the homeodomain necessary for the binding of this transcription factor to its DNA target sites (Figure 1B), due to a protein truncation from the 40th position of the homeodomain onwards, replacing 34 residues by a 10 residues frameshifted sequence (Figure 1C).

**Figure 1.**
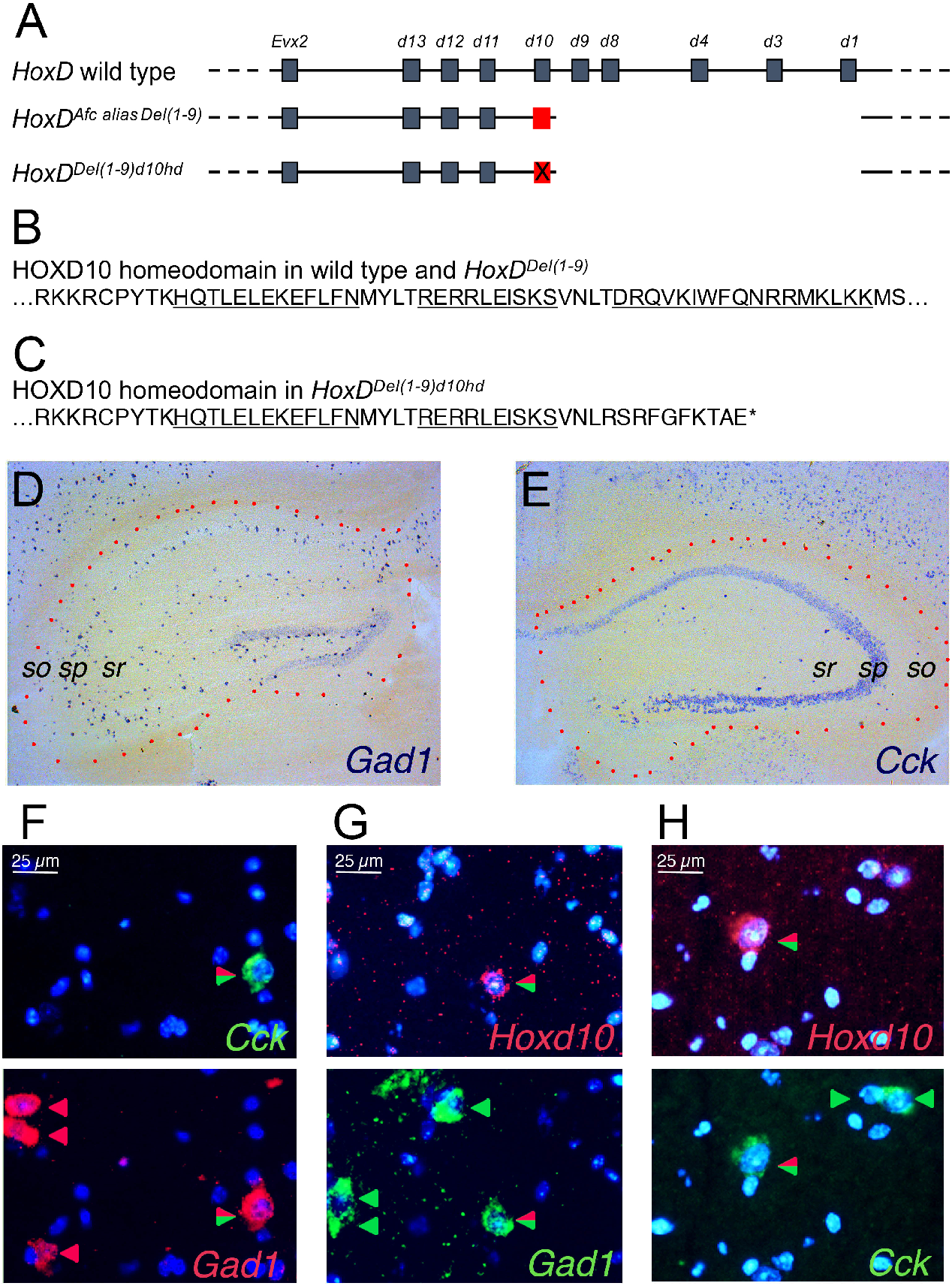
Inactivating mutation in HOXD10 ectopically expressed in a minor GABAergic subpopulation in adult *HoxD^Afc^* brain. (A) Comparison of wild type *HoxD* and the *HoxD^Afc allas Del(1-9)^* and *HoxD^Del(1-9)d10hd^* mutant alleles. Discontinuity of the horizontal line indicates the absence of the genomic segment and the red X indicates the position of the CRISPR/Cas9 hit in the *Hoxd10* homeobox, leading to the generation of the *HoxD^Del(1-9)d10hd^* allele. (B) Amino-acid sequence of the HOXD10 homeodomain in both the wild type and the *HoxD^Afc^* alleles. The three alpha helical subdomains are underlined. (C) Amino-acid sequence of the homeodomain in the truncated HOXD10hd protein product. The sequence of the remaining two alpha helical subdomains are underlined and an asterisk indicates an out of frame stop codon. (D, E). Details of representative coronal sections of heterozygous *HoxD^Afc^* female brains. The contours of the hippocampal formation are indicated by red dots and three landmark cytoarchirectonic layers are annotated (sr, sp, so, for *strata radiatum, pyramidale* and *oriens*, respectively). (D). *Gad1* specific antisense probe reveals positive cells distributed in all layers of CA. (E). A *Cck* specific antisense probe shows few strongly stained cells in all layers of CA, and a relatively weaker signal in the rest of the cells located in *sp*. (F, G, H). Simultaneous fluorescent *in situ* hybridization. Nuclei are shown in blue. (F) Expression of *Cck* in green (top) is detected in one of four *Gad1* positive cells shown below in red (bottom) in CA3 *sr*. (G). Expression of *Hoxd10* (red, top) in one of four *Gad1* positive cells (green, bottom) in CA3 *so*. (H). Expression of *Hoxd10* in (red, top) in one of the three *Cck* positive cells (green, bottom) in CA1 *so*.

We crossed this allele out through three consecutive generations and observed twelve adult females caged with males. Studs were followed for the appearance of injuries at their external genitals. Heterozygous *HoxD^Del(1-9)d10hd^* females bred successfully, without any indications of atypical female courtship (0 out of 12). This was in marked contrast with the observation of 12 out of 18 *HoxD^Afc^* females carrying the intact *Hoxd10* homeobox sequence and showing genital biting [1]. Other abnormal phenotypic traits associated with the *HoxD^Afc^* allele, like malocclusion and slow postnatal weight gain were also rescued [2]. These results provide strong genetic evidence of the direct role of the *HOXD10* transcription factor in bringing about the courtship aberration observed in *HoxD^Afc^* mice.

This courtship anomaly occurred in animals with a low abundance of *Hoxd10* positive cells in adult forebrain, which display molecular and neuroanatomical characteristics reminiscent of a small subpopulation of GABAergic interneurons [1,4], as characterized by the simultaneous detection of both the *Gad1* and *Cck* markers. Double labeling FISH analyses with *Hoxd10-dig* and *Gad1-fluo* pair of probes indeed showed *Hoxd10* positive cells localized selectively in the hippocampus, distributed in any of the layers of CA fields where it co-localized with *Gad1* (Figure 1). Furthermore, by using *Cck-flou* and *Hoxd10-dig* probes, we scored the *Hoxd10* specific red signal in cell accumulating *Cck* transcripts (Figure 1). As all *Cck* positive non-principal cells seemed included in the *Gad1* labeled pool, and since all *Hoxd10* positive cells were part of the *Cck* positive non-principal pool, we concluded that ectopic *Hoxd10* transcripts accumulated in a very sparse subpopulation of *Cck* positive GABAergic cells. Of note, *Hoxd10* like other *Hox* genes is not expressed in any cells of a normal adult forebrain [5].

The *HoxD^Afc^* phenotype followed a gender-specific pattern of expressivity, despite the fact that ectopic expression of *Hoxd10* was similar in both sexes. The ectopic presence of this HOX product in CCK positive GABAergic neurons in adult hippocampus may thus interfere with the implementation of a particular genetic program in a sexually dimorphic manner, perhaps through the property of such proteins to exert a dominant negative effect in various contexts [6]. CKK signaling was previously associated with a sex-dependent control of behavior and its level seems to be modulated during the estrus cycle [7]. Also, the inactivation of the Cck2 receptor, which presumably mediates some effects of CCK neuropeptides in postsynaptic neurons, elicits behavioral alterations distinct in females as compared to males [8]. Altogether, this is consistent with a gender-specific role of CCK positive GABAergic cells in the modulation of behavior [9]. A persistent ectopic expression of *HOXD10* in CCK positive hippocampal GABAergic cells may thus interfere with the function of these cells in controlling the dynamic physiological status of females during the estrous cycle [10].

## Material and Methods

Experiments were conducted according to the Swiss law on animal protection (LPA) under licenses #GE/81/14 and #GE/29/26. The CRISPR/Cas9 induced allele was described in [2]. Freshly dissected brains were mounted in OCT and stored at −80°C. In most experiments, pairs of hemi-brains of *HoxD^Afc^* and *HoxD^Del4-9^*) heterozygous or wild type control adult females were mounted in the same block, cut, collected on the same slides and processed together to allow for direct comparison of the *Hoxd10* signals under identical conditions. Usually four parallel sub-series of 14 *μ*m thick coronal cryo-sections were collected, air-dried and stored at −80°C. One of the sub-series was stained with Cresyl violet and the position of the sections along the Coronal Allen Brain Atlas was determined. On the day of hybridization, slides were thawed, air-dried and fixed in 4% paraformaldehyde in PBS. *In situ* hybridizations were carried out at 63.5°C overnight, followed by stringency washes at 61°C. The binding of the antisense probe was revealed either by the NBT/BCIP alkaline phosphatase substrate (e.g. Allen Brain Institute http://mouse.brain-map.org/gene), or with the FASTRED alkaline phosphatase substrate to detect DIG labeled probes and the Tyramide amplification procedure (PerkinElmer SAT700001EA), followed by Streptavidin Alexa Fluor 488 conjugate (Invitrogene S32354) to detect Fluorescein labeled probes. *Gad1* and *Cck* antisense riboprobes were synthesized using cDNA plasmid clones as templates (http://www.imagenes-bio.de). Briefly, mouse *Gad1* cDNA clone IRAKp961I2154Q was linearized with Kpn1 and transcribed by T7 polymerase (Promega, #P2075). Mouse *Cck* cDNA clone IRAVp968E034D was linearized with EcoRI and transcribed with T3 polymerase (Promega, #P2083). Mouse *Hoxd10* cDNA clone [1] was digested with EcoRI and transcribed by T7 polymerase. Labeled nucleotides were incorporated using DIG RNA Labeling Mix (Roche 1122707390), or Fluorescein RNA Labeling Mix (Roche 11685619910). We successfully detected *Hoxd10* with DIG, yet not when a fluorescein labeled antisense cRNA probe was used. This may reflect a higher sensitivity of the alkaline phosphatase enzymatic reaction, which was also supported by the easier detection of the *Gad1* and *Cck* signals with DIG/FAST RED, as compared to the fluorescein/Tyramide enhancement. In double FISH experiments *Hoxd10* specific red signal was scored at probe concentrations, when red stained cellular profiles were detected only in the *HoxD^Afc^*, and not in either control samples, indicating that conditions were appropriate for specific detection of *Hoxd10* transcripts. The double FISH procedure was carried out as in [11]. Pictures were taken with HBO 100 illumination using the appropriate filter sets to visualize red, green and blue fluorescence signals (set 43, 10 and 49 respectively), on a Zeiss Axioplan 2 microscope (Figure 1C–E). *Hoxd10* red hybridization signals were accepted as positive if the signal could be seen with a 5×/0.25 n.a. 0.17 Zeiss FLUAR objective using filter set 43. Upon higher magnification, a clear cytoplasm signal zone included a negative zone corresponding to the position of a cell nucleus (perikaryon). Images were taken with a Leica DFC300 FX digital color camera. Brightness and contrast were adjusted in Photoshop CS3. Red and blue or green and blue double color images were generated using the HDR2 plug-in.

## Acknowledgements

We thank N. Flores-Ramirez for help with *in situ* hybridizations and I. Rodriguez for discussions. This work was supported by funds from the University of Geneva, the Ecole Polytechnique Fédérale (Lausanne), the Swiss National Research Fund (No. 310030B_138662) and the European Research Council grants System*Hox* (No 232790) and Regul*Hox* (No 588029).

